# RustBEARS: Survival-Conditioned Likelihoods, Uniformization Stochastic Mapping, and Uncertainty Quantification for Historical Biogeography

**DOI:** 10.64898/2026.11.20.768172

**Authors:** Jian He, Jingjing Chen, Wanting Chen, Lei Xie

**Author notes:** Correspondence (J.H.); (L.X.).

## Abstract

Probabilistic discrete-range models, including dispersal–extinction–cladogenesis (DEC), DIVALIKE, BAYAREALIKE, founder-event speciation (+J), spatial modifiers, and time stratification, underpin historical biogeography. As empirical datasets grow, however, BioGeoBEARS workflows face slow multi-area optimization and branch-wise rejection stochastic mapping that fails on rare stepping-stone pathways and scales poorly across posterior trees. Standard fixed-tree pruning likelihoods also retain an absorbing null range without conditioning on extant tip survival, which often drives extinction estimates toward zero and inflates founder-event support, whereas DEC^*^ avoids null absorption only by forbidding single-area extirpation. Boundary Hessian failures and single-tree point estimates also leave parameter, model, phylogenetic, and mapping uncertainties unquantified. Here we present RustBEARS, a Rust engine with BioGeoBEARS-compatible R and Python packages. Across 1,339 configurations from 77 studies, RustBEARS matches BioGeoBEARS log-likelihoods (median |Δln*L*| = 9.95 × 10^−14^) and ancestral probabilities (≤ 1.50 × 10^−14^), accelerates 64- to 256-state optimization 2.8- to 17.3-fold on one core and 4.1- to 36.1-fold on eight cores over the fastest BioGeoBEARS setting, and replaces trial-limited rejection with Uniformization stochastic mapping (170.9-fold median speedup for 200-map Python runs across 30 studies). RustBEARS also introduces whole-tree tip-survival conditioning for unstratified and chronologically emerging island models alongside five frequentist uncertainty tools. Across 58 extant clades from this 77-study archive, survival conditioning cuts near-zero DEC extinction estimates from 62.1% to 0.0% (87.9% finite interior, 12.1% ceiling-censored) and lowers AICc preference for DEC+J from 75.9% to 36.2%.

## Introduction

Reconstructing how lineages expand, contract, and fragment across discrete regions on time-calibrated phylogenies is central to comparative biology (Ronquist 1997; Ree et al. 2005; Ree and Smith 2008; Lamm and Redelings 2009; Ronquist and Sanmartín 2011; Matzke 2013, 2014). The R package BioGeoBEARS (Matzke 2013, 2014) unified classic biogeographic hypotheses by combining branch-wise range expansion (rate *d*) and local extirpation (rate *e*) with sympatric, subset-sympatric, vicariant, and founder-event jump (weight *j*) cladogenesis, enabling likelihood comparisons across DEC, DIVALIKE, and BAYAREALIKE models alongside time stratification and stochastic mapping (Nielsen 2002; Revell 2012; Dupin et al. 2017).

Growing taxon sampling and geographic resolution now strain optimization and pathway sampling in multi-area and multi-tree analyses. With *r* regions, unconstrained state spaces grow as *S* = 2^*r*^ and dense matrix exponentiation scales as *O*(*S*^3^) (Landis et al. 2013; Bettisworth et al. 2023). BioGeoBEARS links interpreted R tree-traversal loops to Fortran 77 matrix exponentials and C++ cladogenetic helpers; cross-language copying, R interpreter overhead, and socket-cluster serialization slow optimization and complicate compilation. Its branch-wise rejection stochastic mapping also simulates forward trajectories repeatedly until matching sampled endpoints, exhausting trial limits on newly emerged islands or rare stepping-stone routes. Graphical tools such as RASP 4 (Yu et al. 2020) broaden multi-model access through BioGeoBEARS and C++ backends, while Lagrange-NG (Bettisworth et al. 2023) accelerated pure DEC point estimation in C++ without +J models, DIVALIKE and BAYAREALIKE cladogenesis, stochastic mapping, BioGeoBEARS R compatibility, or uncertainty propagation (Supplementary Table S1A). Bayesian implementations in RevBayes (Landis et al. 2013; Höhna et al. 2016) are flexible, yet require custom scripts and intensive Markov chain Monte Carlo (MCMC) sampling.

Beyond computation, a longstanding statistical debate concerns why DEC-family models commonly estimate near-zero local extinction 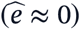 while strongly favoring founder-event speciation (+J) (Lamm and Redelings 2009; Massana et al. 2015; Ree and Sanmartín 2018; Matzke 2022). Along branches, lineages lose areas at rate *e* and enter an absorbing null range (∅) upon losing their final area, yet standard pruning likelihoods do not condition on all sampled extant tips surviving with non-empty ranges. Because raising *e* drains probability into ∅, unconditioned optimization penalizes extinction and compensates via cladogenetic jumps. Massana et al. (2015) proposed DEC^*^ to block null absorption by deleting ∅, preserving multi-area contraction while forbidding single-area endemics from losing their last area. Branch-wise non-null row normalization in RevBayes likewise differs from conditioning on joint extant tip survival across the tree.

Routine biogeographic inference is also limited by how parameter, model, and phylogenetic uncertainties are quantified. Resampling across trees or bootstrap replicates is costly, and boundary effects create statistical difficulties: because *d, e*, and *j* are bounded below by zero, maximum-likelihood estimates often lie on physical boundaries where finite-difference Hessians are singular or indefinite, leaving Wald intervals undefined and breaking *χ*^2^ likelihood-ratio approximations (Self and Liang 1987). Point estimates alone also hide flat high-extinction likelihood ridges in clades dominated by single-area endemics, and BioGeoBEARS lacks integrated tools for multimodel averaging, clade-support-stratified posterior bounds, and stochastic mapping precision checks.

Here we describe RustBEARS, a Rust engine with synchronized R and Python packages addressing these computational and statistical challenges (Fig. 1A; Supplementary Table S1A). RustBEARS retains all 545 BioGeoBEARS R exports, delivers single- and multi-core speedups, and replaces trial-limited rejection with endpoint-conditioned Uniformization stochastic mapping. It also implements fixed-tree tip-survival conditioning across unstratified and chronologically emerging island models alongside five frequentist uncertainty modules. Below we test numerical equivalence across 77 published studies, evaluate statistical behavior across 3,100 simulated datasets and 3,400 mapping trials, reanalyze the 58 eligible extant clades from that archive, and demonstrate workflows in Neotropical *Trimezieae* and Hawaiian *Psychotria*.

**Figure 1.**
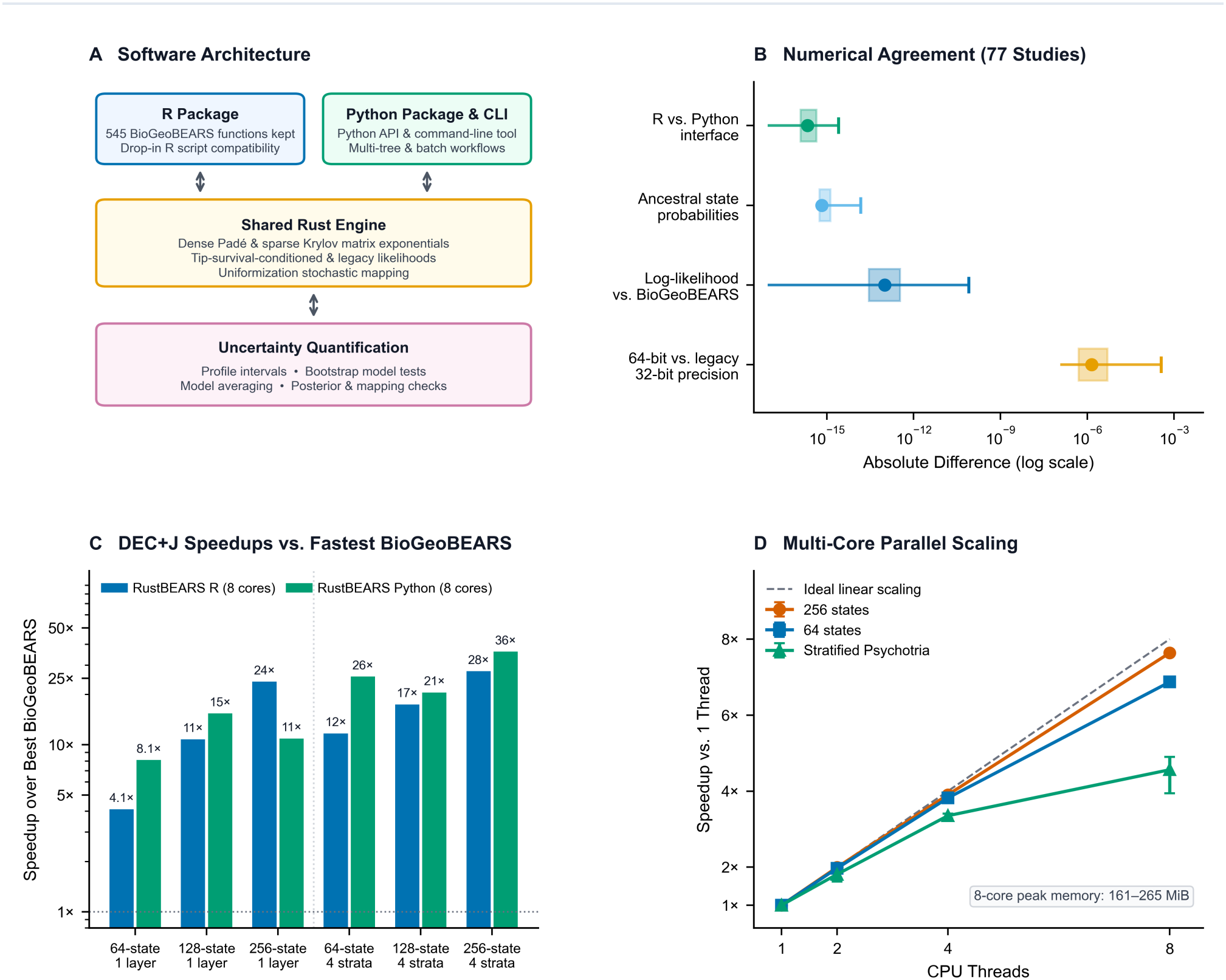
Software architecture, numerical agreement, and computational performance of RustBEARS. **(A)** Overview of the R and Python user packages, the shared Rust computational engine, and the integrated uncertainty quantification modules. **(B)** Absolute numerical differences (median, interquartile box, and fifth-percentile-to-maximum whiskers on a logarithmic scale) across 77 published studies comparing the R and Python interfaces, ancestral state probabilities and log-likelihoods in legacy mode against BioGeoBEARS, and default 64-bit double precision against historical 32-bit mixed precision. **(C)** Eight-core runtime speedups of the RustBEARS R and Python interfaces relative to the fastest one- or eight-core BioGeoBEARS configuration across 64-to 256-state single-layer and four-stratum DEC+J maximum-likelihood optimizations. **(D)** Multi-core strong scaling from one to eight CPU threads (medians and interquartile ranges across five repeats) alongside peak memory usage for full eight-core uncertainty workflows.

## Models and Methods

### Software Architecture and Compatibility with BioGeoBEARS

RustBEARS separates its R and Python interfaces from a memory-safe Rust core built from source with only a standard Rust toolchain, without external Fortran or C++ compilers (Fig. 1A; Supplementary Material S1.1). Branch transition probabilities exp(**Q***t*) use dense Padé scaling-and-squaring on small-to-moderate state spaces and sparse Krylov subspace iterations exp(**Q***t*)**v** (Moler and Van Loan 2003) on larger spaces, alongside sparse coordinate-list cladogenetic kernels. Tree topologies and time-stratified branch segments are precompiled once into contiguous buffers and reused across optimization, profiling, and resampling, avoiding repeated heap allocations and socket serialization.

By default, rate-matrix construction, cladogenetic normalization, matrix exponentials, and log-space pruning use 64-bit IEEE-754 double precision, while an optional legacy mixed-precision mode emulates 32-bit rounding points of upstream C++ helpers so published results can be audited directly. In R, RustBEARS exports all 545 public functions, S4 classes, and plotting utilities of BioGeoBEARS so existing scripts run unchanged once loaded; R and Python also provide concise functions for survival-conditioned and uncertainty analyses (Supplementary Table S1B).

### Survival-Conditioned Likelihoods and Chronological Area Emergence

Over *r* discrete regions with state space *S* = {∅} ∪ *S*_+_ of size *S* = |*S*| (and optional range-size cap *R*_max_ ≤ *r*), range evolution in stratum *m* follows rate matrix **Q**^(*m*)^ , where single-area states {*a*} contract to ∅ at rate 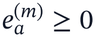. At internal nodes, non-empty ancestral ranges split via normalized cladogenetic probabilities *C*^(*m*)^(*i* → *j, k*) under DEC, DIVALIKE, BAYAREALIKE, or +J models with optional spatial or environmental modifiers.

RustBEARS distinguishes three likelihood formulations on an *N*-tip phylogeny (Supplementary Table S1A; Supplementary Material S1.2). The unconditioned legacy default sets cladogenetic transitions from ∅ to zero and combines root conditional likelihoods **L**_root_ with unnormalized historical root weights without survival conditioning, while disabling ∅ recovers DEC^*^ (Massana et al. 2015). A second, complete unconditional formulation sets *C*^(*m*)^(∅ → ∅, ∅) = 1 under a normalized root prior *π* on *S*_+_, so probabilities sum to unity across all *S*^*N*^ tip outcomes (including extinct tips *Y*_*i*_ = ∅). Under matched branch segmentation and uniform weights across the *S*_0_ allowed non-empty root states (*π*_*i*_ = 1/*S*_0_, with *S*_0_ = *S* − 1 when unconstrained), this log-likelihood on extant non-empty tips differs from unit-weight legacy pruning by −ln*S*_0_ and supports complete-space simulation and probability-sum checks.

Analogous to ascertainment conditioning for discrete characters (Lewis 2001), RustBEARS introduces exact tip-survival conditioning for extant clades. Single-area states still contract into ∅ at rate *e*, and observed tip ranges *Y* = (*Y*_1_, …, *Y*_*N*_) are conditioned on every sampled tip occupying a non-empty present-day range. I an unstratified model with root prior *π* over *S*_+_, a second pruning pass 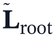 , initialized with ones across allowed present-day non-empty states and zero at ∅, computes exact joint survival 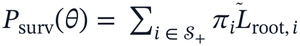 and yields:

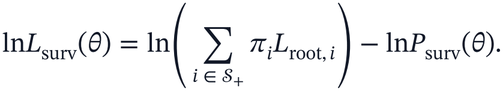

Unlike DEC*, tip-survival conditioning retains single-area extirpation along branches and matches survival-conditioned forward simulation. At fixed parameters, conditioning leaves latent ancestral-state and mapping posteriors unchanged; reconstructions shift because parameter estimates and model weights change.

Tip-survival conditioning extends to spatial, environmental, adjacency, and area-size modifiers and to multi-stratum slicing (0 = *τ*_0_ < *τ*_1_ < … < *τ*_*M*_) with chronologically emerging areas. When areas emerge progressively toward the present (*A*_*M*_ ⊆ *A*_*M* − 1_ ⊆ … ⊆ *A*_1_), every range allowed in older stratum *m* remains valid entering younger stratum 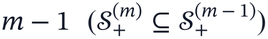. Forward propagation across boundaries thus conserves probability mass without ad hoc renormalization, root summation runs strictly over oldest-stratum non-empty states 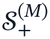 with prior *π*^(*M*)^, and double-precision telescoping sub-branches preserve exact branch-length sums on near-ultrametric trees. Non-monotone masks where occupied areas vanish abruptly (*A*_*m*_ ⊈ *A*_*m* − 1_) lose probability mass without physical extirpation and are rejected outside legacy mode.

### Endpoint-Conditioned Stochastic Mapping via Uniformization

Alongside legacy branch-wise rejection, RustBEARS implements whole-tree Uniformization stochastic mapping. Internal ancestral states and cladogenetic splits are drawn top-down from exact conditional posteriors. For each branch or stratum segment (*a, b*) over Δ*t*, anagenetic jumps and intermediate states are sampled by endpoint-conditioned Uniformization (Hobolth and Stone 2009): **Q**^(*m*)^ is dominated by 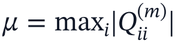, Poisson jump tails are truncated only below a tail-truncation tolerance of 2^−54^, and **P** = **I** + **Q**^(*m*)^/*µ* is filtered through dynamic-programming reachability masks. Uniformization avoids trial-limit failures on rare pathways, respects stratum masks, and yields deterministic histories per map index regardless of thread count.

### Quantifying Parameter, Model, Phylogenetic, and Mapping Uncertainty

Five frequentist uncertainty modules build on these routines (Supplementary Table S1A; Supplementary Material S1.3). Profile likelihood intervals re-optimize nuisance parameters over an adaptive grid, locate two-sided 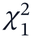 bounds (or legacy support intervals) by Brent root-finding, probe physical zero to separate boundary optima from infeasible zeros, and flag unbounded high-extinction ridges at the search ceiling. Parametric bootstrap LRTs simulate *B* null datasets and refit both models (with null-optimum warm starts for alternatives), calibrating boundary tests where naive resampling or asymptotic tables can fail (Andrews 2000); they report conservative *p*-values (*b* + 1)/(*B* + 1) from *b* exceedance replicates (Boos and Zhang 2000), binomial Monte Carlo envelopes, and deterministic bounds over non-converged replicates. Multimodel inference averages ancestral probabilities and mapping quotas via AIC or AICc weights (*N*-tip convention; Burnham and Anderson 2002). Posterior tree propagation (Nylander et al. 2008; Smith 2009) refits models and maps across external tree sets under fixed sequence-based topology weights, reporting clade monophyly support (*P*_clade_), clade-conditional ancestral probabilities, and deterministic bounds [*P*_min_, *P*_max_] across missing fits (Supplementary Material S4). Stochastic mapping diagnostics compute Monte Carlo standard errors (MCSE) across fixed tree sets or posterior populations, with *t*_*K* − 1_ intervals for *K* independent trees, Geyer’s (1992) initial positive sequence estimator gated at 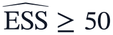 for MCMC chains, and ±*δ* equivalence tests for dispersal asymmetries.

## Results

### Numerical Equivalence and Computational Performance

We tested numerical equivalence across 77 studies comprising 1,339 fixed-parameter configurations (693 unstratified baseline models plus 646 stratified or modifier configurations; 75 valid screens up to 638 states, plus two initial-screen timeouts recovered up to 848 states), 4,888 R assertions, and 243 Python tests (Fig. 1B; Supplementary Table S2A). In legacy mixed-precision mode, RustBEARS matched BioGeoBEARS fixed-parameter log-likelihoods with median |Δln*L*| = 9.95 × 10^−14^ (maximum 8.00 × 10^−11^) and ancestral probability differences ≤ 1.50 × 10^−14^. R and Python interfaces agreed across all 77 studies to median relative |Δln*L*| = 2.13 × 10^−16^ (maximum 2.56 × 10^−15^). Switching to 64-bit precision shifted fixed-parameter log-likelihoods by a median of 1.41 × 10^−6^ (maximum 3.62 × 10^−4^), retained 99.86% (692/693) cross-precision root concordance (98.70% [684/693] relative to BioGeoBEARS) and 100% (693/693) matched-precision optimization non-inferiority across baseline fits, and completed 14 stratified or large-state optimizations where BioGeoBEARS timed out. Across 30 studies (200 maps each), Uniformization matched BioGeoBEARS event proportions closely (median total variation distance 0.0019; mean route *r* = 0.9973, median 0.9995).

On an idle AMD EPYC 7763 server (Fig. 1C; Supplementary Table S2B), RustBEARS accelerated 64-to 256-state unstratified and four-stratum DEC+J optimization 2.8-to 17.3-fold on one core and 4.1-to 27.5-fold in R or 8.1-to 36.1-fold in Python on eight cores over the fastest one- or eight-core BioGeoBEARS run (10.9-to 205.5-fold in strict eight-core comparisons). Socket-cluster overhead made eight-core BioGeoBEARS slower than one core on 64–128 states and on all four-stratum runs (four-stratum 256-state runtime rose from 187.8 s to 261.5 s, whereas unstratified 256-state went from 115.4 s to 99.0 s). Eight-core RustBEARS completed four-stratum 256-state optimization in 5.21 s in Python (36.1-fold over 187.8 s; 50.2-fold over 261.5 s) and 6.83 s in R (27.5-fold; 38.3-fold). For 200-map stochastic mapping, Python Uniformization ran 37.9-to 1,405-fold faster on one core and 25.9-to 212.7-fold faster on eight cores (30-study median: 170.9-fold over single-process BioGeoBEARS). Native R mapping ran 9.8-to 61.2-fold faster on one core for 200 maps (amortizing ∼1.1 s of R table formatting on larger batches) and sampled rare terminal stepping-stone bridges (3.56 × 10^−8^ over 0.25 Ma) where 40,000-trial rejection failed. Across five repeats, eight-thread Python Uniformization scaled 7.64-fold for 256 states (29.70 s to 3.89 s, 1,000 maps), 6.87-fold for 64 states, and 4.56-fold for four-stratum *Psychotria* (5,000 maps); eight-core six-stage workflows used 161–232 MiB of peak RAM for 16–64 states and 168–265 MiB for 256 states (Fig. 1D; Supplementary Table S2C).

### Simulation Validation of Survival Conditioning and Uncertainty Calibration

Across 3,100 simulated datasets and 3,400 mapping trials (Fig. 2; Supplementary Tables S3A–S3B), we first tested null founder-event behavior (*j* = 0) on a 30-tip, four-area (16-state) DEC tree (*d* = 0.015, *e ∈* {0.02, 0.08, 0.15}, 200 datasets per rate). Unconditioned DEC collapsed 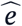 to zero in all 600 datasets (100%), producing an artificial zero-estimate RMSE equal to *e*_true_ (0.020–0.150) and inflating false AICc selection of DEC+J from 5.5% (*e* = 0.02) to 18.0% (*e* = 0.08) and 22.0% (*e* = 0.15). With tip-survival conditioning, median 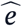 tracked true rates (0.026, 0.093, and 0.145; 1.0–2.5% hit 5.0), although single-tree sampling variance combined zero-boundary mass (23.0–42.5%) with right skewness (means 0.272–0.377; RMSE 0.766–0.945). False AICc selection of DEC+J dropped to 0.5–1.5% (Fig. 2A).

**Figure 2.**
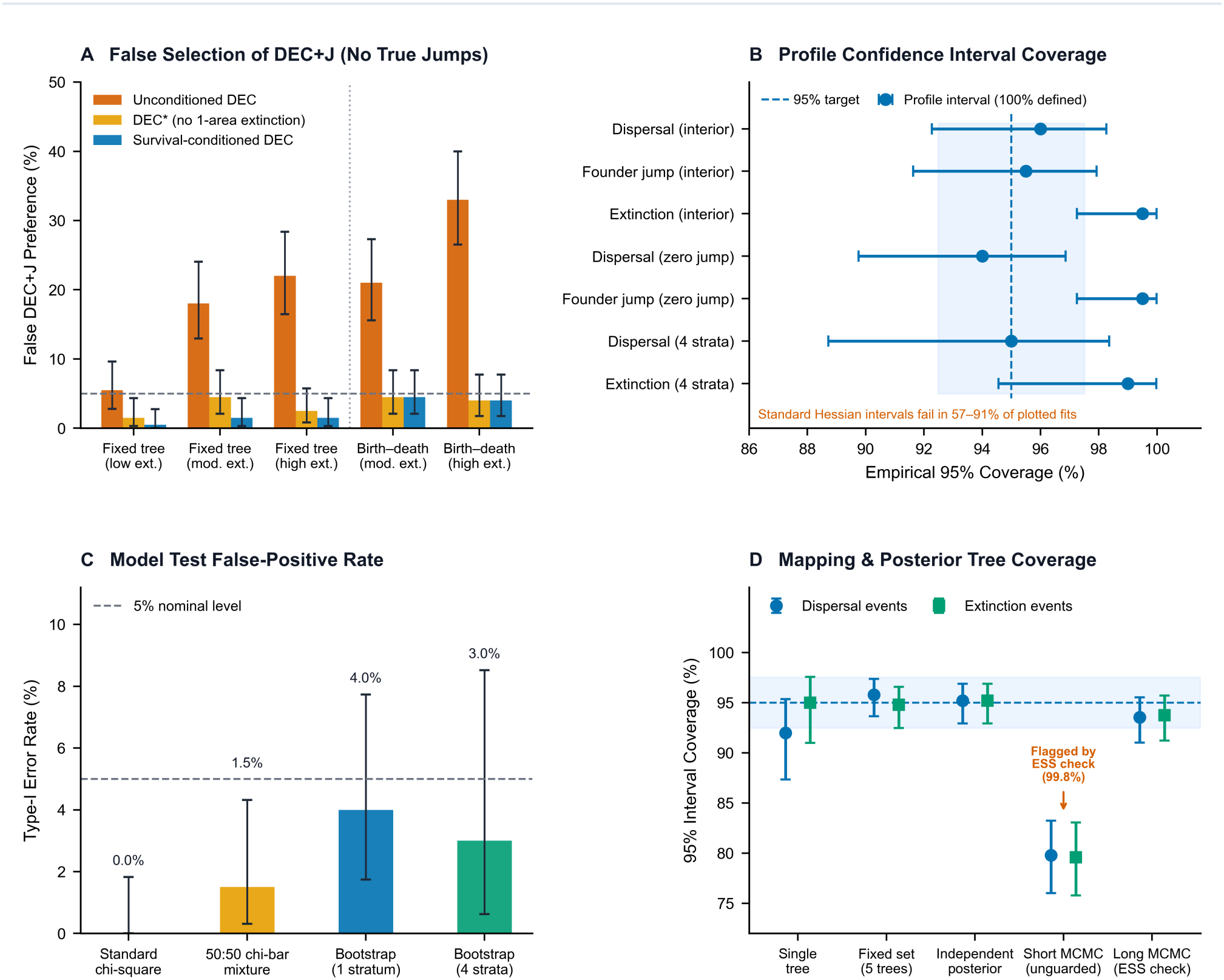
Simulation validation of tip-survival conditioning and calibration of uncertainty methods. **(A)** Percentage of simulated datasets without founder-event speciation in which AICc falsely favored DEC+J (bars and 95% confidence intervals; horizontal dashed line marks 5%) across fixed trees at low, moderate, and high extinction rates and simulated birth–death trees pruned of extinct lineages, comparing unconditioned DEC, DEC*, and survival-conditioned DEC. **(B)** Empirical 95% coverage of profile likelihood intervals (points and 95% intervals; vertical dashed line and shaded band mark the 95% target) across unstratified interior, zero-boundary, and four-stratum island-emergence simulations, where standard Hessian intervals fail in 57–91% of fits. **(C)** False-positive (Type-I error) rates at the 5% significance level (bars and 95% intervals) when testing survival-conditioned DEC against DEC+J under standard chi-square, 50:50 chi-bar mixture, and parametric bootstrap likelihood-ratio tests across single-stratum and four-stratum models. **(D)** Empirical 95% coverage of stochastic mapping event intervals (points and 95% intervals) across single trees, five-tree sets, independent posterior tree samples, unguarded short MCMC chains, and long MCMC chains passing the effective-sample-size check.

To check whether this result depended on a fixed tree, we simulated 400 independent 30-tip phylogenies (plus 200 DEC*-truth control trees) under a forward birth–death and DEC process (first 30-lineage sojourn with* ≥ 2 *occupied areas; Hartmann et al. 2010) at λ* = 0.30, *d* = 0.03, *e* = 0.08 *or λ* = 0.35, *d* = 0.04, *e* = 0.15, *where losing the final area caused extinction (j* = 0, *200 trees per regime; Fig. 2A). Unconditioned DEC collapsed* 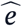 *to zero on 84*.*5–90*.*0% of trees and falsely selected DEC+J by AICc on 21*.*0% (e* = 0.08*) and 33*.*0% (e* = 0.15*). Tip-survival conditioning recovered median extinction closely (0*.*079 for e* = 0.08; *0*.*183 for e* = 0.15, *lowering median absolute error from 0*.*150 to 0*.*111; 0*.*101 on the e* = 0.10 *control), while single-tree right-tail dispersion persisted (extinction RMSE 0*.*575 and 0*.*323; dispersal RMSE 0*.*011 and 0*.*017, matching unconditioned 0*.*011 and 0*.*017). It cut AICc false positives to 4*.*5% and 4*.*0% (0*.*0–1*.*0% ceiling-censored, matching DEC*) and yielded lower high-extinction RMSE than DEC^*^ for extinction (0.323 versus 0.423) and dispersal (0.017 versus 0.027).

Across 800 four-area datasets spanning four interior and boundary regimes, central-difference Hessian prechecks failed at box boundaries in 54–91% of fits (57–91% in Fig. 2B). By contrast, 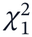 profile intervals were defined in 100% of fits, with 94.0–98.5% coverage on interior *d* and *j* and 99.5–100.0% at zero boundaries and on *e* (right-censored at 5.0 in 18.5–83.0% of 40-tip datasets). In four-stratum island-emergence simulations (30 tips, 100 datasets; true *d* = 0.08, *e* = 0.06; median 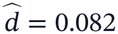,ê = 0.028), stratified profile intervals achieved 95.0% coverage on *d* and 99.0% on *e* with zero ceiling censoring. Boundary point masses made asymptotic 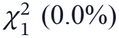 and 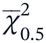 (0.0–1.5%) tests conservative under survival conditioning, whereas parametric bootstrap LRTs (*B* = 99) held Type-I error near 5% in unstratified (4.0%) and four-stratum (3.0%) models (Fig. 2C; power at *j* = 0.03 was 11.0% versus 1.5% for *χ*^2^). Across 600 multimodel datasets with comparable candidate weights, Akaike averaging cut ancestral Brier error by 0.048 and 0.022 relative to single-best-model selection (*p* < 10^−6^) and avoided +∞ log-loss (up to 21.0% of single-model fits). In 3,400 mapping trials (Fig. 2D), single-tree, five-tree fixed-set, and *t*_9_ independent-tree coverage was 92.0–95.8%. On autocorrelated MCMC chains (*ρ* = 0.8), the 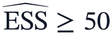 check caught 99.8% (499/500) of short chains (*K* = 60, true effective trees = 8.4, unguarded coverage 79.6–79.8%) and passed 99.0–99.2% of long chains (*K* = 800, median 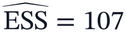) at 93.5–93.7% conditional coverage.

### Empirical Reassessment Across 58 Archived Clades

Comparing unconditioned and tip-survival-conditioned fits across the 77-study archive under standardized DEC and DEC+J bounds, we excluded 10 fossil-tip entries (≥ 1 Ma), four non-ultrametric extant trees (0.004–0.401 Ma), one survival-underflow case, two entries above 512 states, and two timeout baselines, leaving 58 extant clades (Fig. 3A; Supplementary Table S3C). Under unconditioned DEC, 62.1% (36/58) estimated near-zero extinction 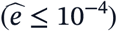. Under tip-survival conditioning, near-zero DEC estimates fell to 0.0% (minimum 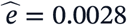), and 87.9% (51/58) yielded finite interior estimates below 5.0 (interior median 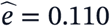; overall median 0.186). The remaining 12.1% (7/58) of DEC fits (5.2% [3/58] in DEC+J) hit 5.0 with tiny survival probabilities (ln*P*_surv_ = −325.5 to −6, 419.3), reflecting flat upper-tail likelihood ridges in single-area-dominated clades rather than precise extreme rates; this group included all four (100%) datasets where every tip occupied one area (censored median single-area fraction 100.0%, range 70.0–100.0%, versus 83.3% among interior entries). Paired AICc preference for DEC+J dropped from 75.9% (44/58; median *D* = 18.70) to 36.2% (21/58; median *D* = 0.64), with 23 clades shifting to DEC and none reversing; excluding the seven ceiling-censored entries yielded a similar drop from 72.5% (37/51) to 39.2% (20/51).

**Figure 3.**
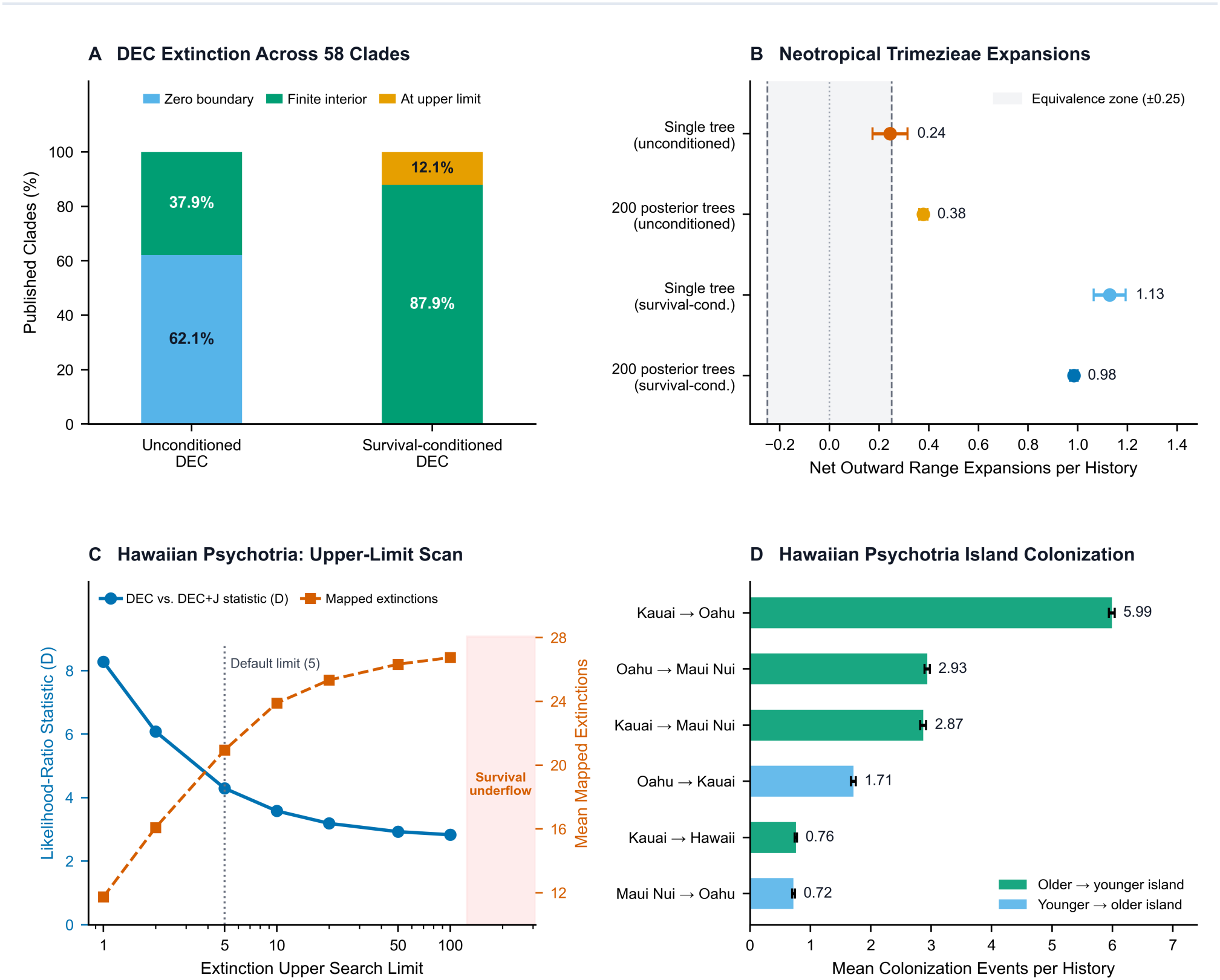
Empirical reassessment of 58 published clades and case studies in Neotropical *Trimezieae* and Hawaiian *Psychotria*. **(A)** Proportions of published clades yielding zero-boundary, finite interior, or upper-limit extinction estimates under unconditioned and survival-conditioned DEC. **(B)** Net outward minus inward range expansions per mapped history from *campo rupestre* to surrounding habitats in Neotropical *Trimezieae* (Vasconcelos et al. 2020) on the reference tree and across 200 posterior trees (points and 95% Monte Carlo intervals; shaded band marks the ±0.25 equivalence zone). **(C)** Sensitivity of the DEC versus DEC+J likelihood-ratio statistic (blue solid line, left axis) and mean mapped DEC extinction counts (vermilion dashed line, right axis) across increasing upper search limits on extinction in four-stratum Hawaiian *Psychotria*, with the vertical dotted line marking the default limit and pink shading marking numerical survival underflow at extreme rates. **(D)** Model-averaged inter-island colonization events per history (bars and 95% Monte Carlo intervals) across four-stratum Hawaiian *Psychotria*, distinguishing dispersal from older to younger islands (green) from dispersal toward older islands (light blue).

### Empirical Case Studies: Neotropical Trimezieae and Hawaiian Psychotria

We analyzed Neotropical *Trimezieae* (Vasconcelos et al. 2020; 57 terminal lineages across two habitat areas: E, recorded in *campo rupestre*, and N, outside, giving four states) across 200 posterior trees, six models, and 100,000 maps (Fig. 3B). Extrapolated from matched server runs (two trees in 111.75 s for single-core BioGeoBEARS versus 20 trees in 75.07 s for eight-thread RustBEARS), 200-tree runtime fell from ∼3.1 h to 12.5 min (∼1.9-fold per nominal core on four states; retaining 10,000 histories used 1,703 MiB RAM). Across all 200 trees, *d* and *e* had finite interior MLEs whereas +J models estimated 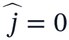 (reference-tree *B* = 399 bootstrap *p* = 0.63–1.00), and BAYAREALIKE was favored on 199/200 trees. On the reference tree, survival conditioning raised BAYAREALIKE extinction from 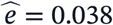 (2-unit support interval [0.011, 0.082]) to 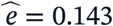 (95% profile CI [0.045, 0.348]; DEC rose from 0.000 to 0.159 [0.000, 0.676], matching 0.161 in DEC*). Mean outward minus inward expansion (E → N) − (N →E) (*δ* = 0.25, conditional on per-tree MLEs) shifted on the reference tree (5,000 maps) from an inconclusive 0.244 ([0.173, 0.314]) to 1.128 ([1.064, 1.193]). Across 200 posterior trees (100,000 maps), it rose from 0.378 ([0.361, 0.395]) to 0.984 ([0.969, 1.000]; relative MCSE = 0.80%), tracking monophyly across 25 non-monophyletic clades (zero missing fits among 1,200 optimizations).

In four-stratum Hawaiian *Psychotria* (Nepokroeff et al. 2003; Matzke 2014; 19 single-island-coded terminal lineages across 11 species under chronological emergence from Kauai to Oahu, Maui Nui, and Hawaii; Fig. 3C–D), survival conditioning at ceiling 5.0 lowered the DEC versus DEC+J statistic from *D* = 17.85 to *D* = 4.29, leaving founder-event support borderline (*B* = 999 bootstrap *p* = 0.044, 95% MC envelope [0.031, 0.058]). Profile intervals flagged extinction as right-censored (*e* ≥ 2.05 in DEC; *e* ≥ 1.24 in DEC+J; DEC^*^ MLEs also hit 5.0). Scanning ceilings from 1 to 200 (Fig. 3C) showed why: because all 19 tips are single-island endemics, raising *e* flattens the conditional likelihood (*D* fell from 8.28 to 2.83 at ceiling 100, non-+J Akaike weight reached 50.7%, and underflow was guarded at ≥ 150) as mean mapped DEC extinctions rose from 11.7 to 26.7. Across 5,000 six-model-averaged Uniformization histories at ceiling 5.0 (3.19 s locally; single-model 5,000-map run in 2.16 s on an eight-thread AMD EPYC 7763; zero prohibited transitions), progressive older-to-younger colonization dominated (mean MCSE: ± K→O, 5.99 ± 0.025 ; O→M, 2.93 ± 0.022; K→M, 2.87 ± 0.025; relative MCSE < 1.8%; Fig. 3D). That pattern held across ceilings 1–20 (78.4–88.3% older-to-younger flux with K →O first), although secondary O →M and K → M rankings swapped at ceiling 20 (Supplementary Table S3C).

## Discussion

Modern biogeographic datasets increasingly span many regions and hundreds of posterior trees, placing heavy demands on likelihood evaluation and stochastic mapping. Our implementation builds on Ronquist (1997), Ree et al. (2005), Ree and Smith (2008), Ronquist and Sanmartín (2011), Matzke (2013, 2014), Dupin et al. (2017), and Yu et al. (2020). As state spaces expand, dense matrix exponentiation grows cubically (Landis et al. 2013), while legacy workflows such as BioGeoBEARS (v1.1.4) incur additional overhead passing data among R, Fortran, and C++, and branch-wise rejection mapping (Nielsen 2002; Revell 2012; Dupin et al. 2017) stalls on rare branch endpoints. Following the compiled-language approach demonstrated for pure DEC in Lagrange-NG (Bettisworth et al. 2023), RustBEARS runs the full BioGeoBEARS model set in parallel Rust memory with sparse matrix methods (Moler and Van Loan 2003), and adapts endpoint-conditioned Uniformization (Hobolth and Stone 2009) to sample histories directly between known branch endpoints. Across 64-to 256-state benchmarks, optimization runs 2.8-to 17.3-fold faster on one core and 4.1-to 36.1-fold faster on eight cores over the fastest BioGeoBEARS configuration, while 200-map Python stochastic mapping runs a median of 170.9-fold faster across 30 studies, reducing analyses that took hours or failed on rare routes to seconds or minutes. RustBEARS also preserves full backward compatibility with BioGeoBEARS, retaining all 545 R exports and reproducing published log-likelihoods across 77 studies (median |Δln*L*| = 9.95 × 10^−14^) alongside a synchronized Python interface.

Why DEC models often estimate zero extinction while strongly favoring founder-event speciation (+J) has been debated for over a decade (Matzke 2014, 2022; Massana et al. 2015; Ree and Sanmartín 2018). Lamm and Redelings (2009) identified the survivorship problem: phylogenies of living species include only lineages surviving in at least one region, yet standard likelihoods penalize positive extinction whenever probability flows into the null range. Massana et al. (2015) avoided that penalty in DEC^*^ by removing the null range, assuming single-area species never go extinct. RustBEARS retains single-area extinction, computes exact joint tip survival across unstratified and emerging-island models, and divides the likelihood by that probability (paralleling ascertainment conditioning; Lewis 2001). Simulations clarify both strengths and limits: when *j* = 0, unconditioned DEC pushes extinction to zero and falsely selects DEC+J in 5.5–33% of datasets (18–33% at moderate-to-high extinction), whereas survival conditioning restores median extinction near true values and cuts false DEC+J selection to 0.5–4.5%, although single-tree extinction estimates retain right-skewed dispersion on 30-tip trees. Across the 58 extant clades in our 77-study archive, survival conditioning removes near-zero DEC extinction estimates (62.1% to 0.0%) and lowers AICc preference for DEC+J from 75.9% to 36.2% (72.5% to 39.2% among the 51 non-censored clades). Thus, over half (23 of 44) of previously favored +J cases disappear once zero-extinction bias is corrected (consistent with Massana et al. 2015; Ree and Sanmartín 2018), yet founder-event speciation remains favored in over one-third of clades (consistent with Matzke 2014, 2022).

Biogeographic conclusions can also become overconfident when studies rely on single-tree, single-model point estimates near parameter boundaries. Ignoring phylogenetic uncertainty (Nylander et al. 2008; Smith 2009; Vasconcelos et al. 2020), conditioning on a single best-fit model (Burnham and Anderson 2002), or applying asymptotic formulas at zero boundaries (Self and Liang 1987; Andrews 2000) can understate true uncertainty. Although 87.9% of archived clades yielded a finite extinction estimate below the upper limit, a single point estimate does not guarantee tight constraint: where every species occupies a single island or region (all four 100%-single-area archival datasets and Hawaiian *Psychotria*), moderate and very high extinction rates explain extant tips almost equally well. RustBEARS makes these checks routine via five tools: profile likelihoods trace parameter ranges and flag unidentifiable upper bounds; parametric bootstrap tests (Boos and Zhang 2000) replace invalid *χ*^2^ tables; Akaike weights average reconstructions across models; posterior tree propagation summarizes clade monophyly and missing-fit bounds; and mapping diagnostics check MCSE and MCMC effective sample sizes (Geyer 1992). In Hawaiian *Psychotria*, these checks caution against overinterpreting secondary route rankings that shift with the extinction ceiling, while remaining consistent with progressive older-to-younger colonization (Nepokroeff et al. 2003; Matzke 2014).

Nevertheless, RustBEARS has clear scope boundaries. Although single-area extirpation into the null range induces range-dependent lineage loss along branches, RustBEARS conditions on a fixed time-calibrated tree rather than modeling a joint tree-and-range diversification likelihood. Studies where geographic range directly drives speciation and extinction rates (as in GeoSSE and hidden-state models; Goldberg et al. 2011; Caetano et al. 2018; Matzke 2022) or where phylogeny, fossils, and geography are co-estimated from sequence data (Landis et al. 2013; Matzke 2013; Höhna et al. 2016) should therefore pair RustBEARS with those frameworks. Fossil-tip trees and island-submergence masks remain supported in legacy mode, and extending survival conditioning to fossil lineages and submerging islands is a natural next step.

## Software and Data Availability

RustBEARS (v1.2.0; Rust workspace, R package RustBEARS, Python package and CLI rustbears, user documentation, runnable tutorials, and reproduction benchmark suites) is open-source under the GNU General Public License v3.0 or later at https://github.com/Jhe1004/RustBEARS (release archive v1.2.0: https://github.com/Jhe1004/RustBEARS/releases/tag/v1.2.0). Users publishing analyses with RustBEARS are asked to cite Matzke (2013, 2014) alongside this article.

## Supplementary Material

Online Supplementary Material (Supplementary Tables S1A–S1B, S2A–S2C, and S3A–S3C, mathematical derivations S1.1–S1.3, and external Bayesian posterior tree-propagation analyses S4) accompanies this preprint as a separate PDF file.

## Acknowledgments

Artificial intelligence tools were used to assist with software coding and to polish the language of the manuscript. All code, analyses, and text were reviewed and verified by the authors, who take full responsibility for the content of this work.

