## Supplementary Material for "RustBEARS: Survival-Conditioned Likelihoods, Uniformization Stochastic Mapping, and Uncertainty Quantification for Historical Biogeography"

#### Supplementary Table S1A. Comparison of Probabilistic Historical Biogeography Software and Summary of Analytical Methods Implemented in RustBEARS

##### Panel A. Capabilities Across Probabilistic Historical Biogeography Software

| Analytical Capability | BioGeoBEARS (R; Matzke 2013, 2014) | Lagrange-NG (C++; Bettisworth et al. 2023) | RevBayes (C++; Landis et al. 2013; Höhna et al. 2016) | RustBEARS (v1.2.0; Rust engine with R and Python interfaces) |
|---|---|---|---|---|
| **User interfaces & script compatibility** | R package (545 exported functions & classes) | Command-line interface | Rev probabilistic programming language | **R package retaining all 545 upstream exports (reusable upon loading RustBEARS) + Python API & CLI** |
| **Cladogenetic range-inheritance models** | DEC, DIVALIKE, BAYAREALIKE, and +J variants | DEC only | DEC, +J, and state-dependent diversification (ClaSSE / GeoSSE) | **DEC, DIVALIKE, BAYAREALIKE, and +J variants** |
| **Spatial modifiers & geological stratification** | Distance, environment, area size, adjacency, time slicing, and allowed-area masks | Time-stratified DEC rate matrices and connectivity multipliers | Epoch-stratified rate matrices and paleogeographic connectivity | **All spatial modifiers + multi-stratum time slicing + chronological area emergence** |
| **Extinction & survival conditioning** | Unconditioned default; or DEC* (removes null state, forbidding single-area extinction) | Unconditioned DEC | Branch-wise non-null row normalization (CondSurv) or birth–death diversification | **Whole-tree joint tip-survival conditioning (retaining single-area extinction), complete unconditional, legacy, and DEC*** |
| **Biogeographical stochastic mapping** | Branch-wise rejection sampling (vulnerable to trial-limit failure on rare transitions) | Marginal ancestral state probabilities only | Endpoint-conditioned MCMC or rejection character histories | **Whole-tree endpoint-conditioned Uniformization bridges (eliminating rejection trial-limit failures; thread-deterministic) + legacy rejection** |
| **Uncertainty & multi-tree inference** | Local Hessian SEs; asymptotic $\chi^2$ LRT; AIC/AICc; per-tree fits & master-tree node averaging | Maximum-likelihood point estimates (single-tree & multi-tree batch processing) | Full Bayesian MCMC posterior distributions | **Profile likelihood CIs, parametric bootstrap LRT, Akaike model averaging, clade-support-stratified posterior bounds, and mapping MCSE / ESS diagnostics** |

##### Panel B. Likelihood Conditioning Formulations and Uncertainty Quantification Methods in RustBEARS

| Analytical Component | Target Estimand & Scope | Core Statistical Formulation | Built-in Identifiability & Numerical Diagnostics |
|---|---|---|---|
| **Tip-survival conditioned likelihood** | Conditional probability $P_\theta(Y \mid \forall k,\ Y_k \neq \varnothing)$ on a fixed reconstructed phylogeny | Retains rate-$e$ extirpation from single-area states into $\varnothing$; divides joint tip probability by exact extant survival probability $P_{\mathrm{surv}}(\theta)$ across single or chronologically emerging strata | Exact duality with forward survival-conditioned simulation; guards against IEEE-754 double-precision survival underflow at extreme extinction rates; rejects probability-leaking submergence masks |
| **Complete unconditional distribution** | Joint distribution $P_\theta(Y)$ over all $S^N$ tip outcomes (including extinct tips $Y_k = \varnothing$) | Completes cladogenesis at the null state via $C(\varnothing \to \varnothing, \varnothing) = 1$ under a normalized root prior; under matched branch segmentation and uniform weights over $S_0$ allowed non-null root states, differs from the unit-weight legacy score on extant tips by $-\ln S_0$ ($S_0 = S - 1$ when unconstrained) | Serves as a probability-conserving forward simulation and exact state-space normalization baseline |
| **Unconditioned legacy likelihood** | Historical BioGeoBEARS pruning score and raw root-weight scaling | Retains upstream unnormalized root weights and all historical model extensions (DEC*, trait-coupled dispersal, fossil branches, fixed internal nodes) | Reproduces published results to machine precision in 32-bit mixed-precision mode; requires an explicit generating model prior to parametric simulation |
| **Profile likelihood intervals** | Confidence intervals (conditioned models) or support intervals (legacy score) | Re-optimizes nuisance parameters across an adaptive grid and locates two-sided $\chi^2_1$ bounds via Brent root-finding (with optional local bootstrap or one-sided boundary cutoffs) | Probes exact physical zero to distinguish boundary optima from structurally infeasible zeros; flags unbounded high-extinction likelihood ridges at the optimization ceiling |
| **Parametric bootstrap LRT** | Finite-sample null distribution of $D = 2(\ln L_1 - \ln L_0)$ for nested hypotheses | Simulates $B$ datasets from the fitted null model and refits both models (initializing alternative fits at the null solution); reports conservative $p = (b+1)/(B+1)$ from $b$ exceedance replicates | Retains non-converged replicates in the planned denominator with deterministic failure bounds and 95% Clopper–Pearson Monte Carlo envelopes |
| **Multimodel & posterior tree propagation** | Akaike-weighted ancestral probabilities and stochastic maps across models and tree sets | Combines compatible candidate models via AIC / AICc weights ($N$-tip convention) and propagates inference across external posterior tree samples | Isolates incomplete trees; reports per-clade topological support ($P_{\mathrm{clade}}$) alongside clade-conditional ancestral probabilities and deterministic bounds $[P_{\min}, P_{\max}]$ over any missing tree fits |
| **Stochastic mapping precision & stopping** | Monte Carlo standard errors (MCSE and relative MCSE) of event counts and dispersal routes | Separates fixed tree-sample precision from posterior tree-population extrapolation ($t_{K-1}$ for $K$ independent trees; Geyer [1992] initial positive sequence variance for MCMC chains) | Enforces a calibrated effective sample size threshold ($\widehat{\mathrm{ESS}} \geq 50$) on MCMC tree chains; classifies pairwise route asymmetries via equivalence testing around $\pm\delta$ |

#### Supplementary Table S1B. User-Facing Functions and Minimal Workflow Examples in R and Python

*(All user-facing function names, options, and minimal scripting examples across R and Python are consolidated here for practitioner reference; step-by-step guides are hosted at https://github.com/Jhe1004/RustBEARS.)*

| Analytical Task | R Package ( `library(RustBEARS)` ) | Python Package ( `import rustbears as rb` ) | Key Options & Notes |
|---|---|---|---|
| **Model specification & likelihood conditioning** | `run <- define_BioGeoBEARS_run(...)` `run$likelihood_conditioning <- "tip_survival"` `run$root_prior <- define_root_prior("uniform_nonnull")` | `mod = rb.BioGeoBEARS(tree, geog, model="DEC", likelihood_conditioning="tip_survival", root_prior=rb.RootPriorSpec("uniform_nonnull"))` | Conditioning choices: `"tip_survival"`, `"unconditional_complete"` (with `allow_null_tips`), or `"legacy_unconditioned"` (default) |
| **Precision mode selection** | `options(RustBEARS.precision = "StrictF64")` *(default)* or `"LegacyF32"` | `precision="strict_f64"` *(default)* or `"legacy_f32"` | `"LegacyF32"` emulates 32-bit mixed rounding in upstream `cladoRcpp` |
| **Maximum-likelihood optimization** | `fit <- bears_optim_run(run)` | `fit = mod.fit(num_threads=8)` | Supports `timeperiods`, monotone `areas_allowed`, and `+x/+n/+w/+u` modifiers |
| **Forward dataset simulation** | `simulate_datasets(fit, n_reps = 200L, seed = 42L)` | `rb.simulate_datasets(mod, params=..., n_reps=200, seed=42)` | Paired directly with `"tip_survival"` or `"unconditional_complete"` |
| **Profile likelihood intervals & surfaces** | `profile_ci(fit, params = c("d", "e"), num_threads = 8L)` | `fit.profile_ci(params=["d", "e"], num_threads=8)` `fit_decj.profile_surface("d", "j")` | Probes exact physical zero and flags ceiling ridges via `status_upper == "censored_at_box_upper"` |
| **Parametric bootstrap LRT** | `bootstrap_lrt(fit_dec, fit_decj, n_boot = 399L, checkpoint_file = "lrt.rds")` | `rb.bootstrap_lrt(fit_dec, fit_decj, n_boot=399, checkpoint_file="lrt.jsonl")` | Supports SHA-256 verified `resume=True` checkpoints |
| **Multimodel Akaike averaging** | `model_average(list(fit_dec, fit_decj), criterion = "AICc", num_maps = 2000L)` | `avg = rb.model_average([fit_dec, fit_decj], criterion="AICc")` `maps = avg.run_bsm(num_maps=2000)` | Verifies matching data, conditioning, and root priors |
| **Posterior tree-set propagation** | `fit_over_trees(run, trees, models = c("DEC", "DEC+J"), likelihood_conditioning = "tip_survival", bsm_maps_per_tree = 500L)` | `rb.fit_over_trees(trees, geog, models=[...], likelihood_conditioning="tip_survival", bsm_maps_per_tree=500)` | Pass `likelihood_conditioning` explicitly; reports per-clade monophyly support ($P_{\text{clade}}$) and `[P_min, P_max]` bounds over missing tree fits |
| **Whole-tree Uniformization BSM & diagnostics** | `maps <- runBSM(fit, NULL, nummaps_goal = 2000L, bsm_engine = "native_whole_tree", num_threads = 8L)` `bsm_diagnostics(maps, tree_target = "fixed_sample")` | `maps = fit.run_bsm(num_maps=2000, num_threads=8)` `fit.run_bsm_adaptive(target_rmcse=0.05)` `rb.bsm_diagnostics(maps)` | Enforces $\widehat{\text{ESS}} \geq 50$ for `tree_target="posterior_population"` on MCMC chains; `"legacy_r_bridge"` reproduces upstream rejection BSM |

### S1. Mathematical Formulation, Software Architecture, and Stratified Conditioning

#### S1.1 Shared Rust Crate Architecture and Dual Precision Modes

`RustBEARS` (v1.2.0) separates user-facing R and Python interfaces from a memory-safe Rust workspace (built from source with only a standard Rust toolchain and zero external Fortran or C++ compiler dependencies) comprising six crates:

1. **`bgb-core`**: Canonical discrete geographic state-space indexing (ordered by range cardinality and lexicographic area combinations, preserving exact `BioGeoBEARS` state indices), Newick tree topology compilation, and geological time-stratum slicing.
2. **`bgb-numerics`**: Dense Padé scaling-and-squaring matrix exponentials ($\exp(\mathbf{Q}t)$) for small-to-moderate state spaces and sparse Krylov subspace (`EXPOKIT`-compatible Arnoldi) action $\exp(\mathbf{Q}t)\mathbf{v}$ for high-dimensional state spaces, plus keyed deterministic `Xoshiro256PlusPlus` random number streams.
3. **`bgb-clado`**: Exact sparse coordinate-list (COO) cladogenetic transition kernels supporting sympatric (`y`), subset-sympatric (`s`), vicariant (`v`), and founder-event jump (`j`) speciation weights, along with spatial distance (`+x`), environmental distance (`+n`), manual dispersal multiplier (`+w`), and area-size (`+u` / `+e`) modifiers.
4. **`bgb-step`**: Whole-tree pruning (`downpass`) and ancestral marginal state (`uppass`) propagation across single-layer and multi-stratum phylogenies, precompiling tree topology buffers once per optimization or bootstrap run to eliminate repeated R heap allocations.
5. **`bgb-bsm`**: Dual stochastic character mapping engines: whole-tree endpoint-conditioned Uniformization Markov bridges with dynamic-programming reachability masks, alongside a legacy branch-wise rejection engine (`legacy_r_bridge`) for historical script reproduction.
6. **`bgb-ffi`**: Thread-safe C ABI bindings (`ctypes` in Python and `.Call` in R) enforcing `OPENBLAS_NUM_THREADS = 1` inside multi-worker pools to prevent nested thread oversubscription.

Two arithmetic modes are exposed across both frontends: - **`StrictF64` (default in v1.2.0)**: Full IEEE-754 binary64 arithmetic across rate-matrix construction, cladogenetic weight normalization, matrix exponentiation, and log-space likelihood accumulation. - **`LegacyF32` (`options(RustBEARS.precision = "LegacyF32")` in R / `precision = "legacy_f32"` in Python)**: Reproduces the exact mixed-precision rounding points of upstream `cladoRcpp` C++ routines (32-bit `float` storage/products paired with 64-bit `double` accumulators and decimal step-boundary rounding in `EXPOKIT` stepping) for historical auditability.

---

#### S1.2 Explicit Likelihood Conditioning Conventions and Chronologically Monotone Area Emergence

Let $\mathcal{S} = \{\varnothing\} \cup \mathcal{S}_+$ denote the discrete geographic state space over $r$ regions (with maximum range size $R_{\max} \le r$ and total state count $S = |\mathcal{S}|$), where state 0 represents the null range $\varnothing$ and $\mathcal{S}_+ = \{1, \ldots, S-1\}$ denotes non-empty ranges. Along branches in geological stratum $m$, ranges evolve according to continuous-time Markov rate matrix $\mathbf{Q}^{(m)}$ in which single-area states $\{a\}$ transition to $\varnothing$ at local extinction rate $e_a^{(m)} \ge 0$, while $\varnothing$ is absorbing ($Q^{(m)}_{\varnothing,s} = 0\ \forall s$).

At internal speciation nodes in stratum $m$, let $C^{(m)}(i \to j, k)$ denote normalized cladogenetic inheritance probabilities for non-empty ancestor $i \in \mathcal{S}_+$. `RustBEARS` formalizes three explicit likelihood conventions:

1. **`legacy_unconditioned` (historical compatibility default)**: Sets $C^{(m)}(\varnothing \to \cdot, \cdot) = 0$, evaluates conditional likelihood vectors $\mathbf{L}_{\text{root}}$ from observed tip states $Y = (Y_1, \ldots, Y_N)$, and reports the unnormalized inner product $\sum_{i \in \mathcal{S}_+} w_i L_{\text{root},i}$ (using upstream flat or equilibrium-like root weights $w_i$ without dividing by tip survival). When combined with `include_null_range = FALSE`, this recovers the `DEC*` family (Massana et al. 2015), which removes $\varnothing$ from $\mathcal{S}$ and structurally forces single-area extinction rates to zero.

2. `unconditional_complete` : Completes the cladogenetic kernel on $\varnothing$ via deterministic empty-to-empty transmission $C^{(m)}(\varnothing \to \varnothing, \varnothing) = 1$ with normalized root prior $\pi$ on $\mathcal{S}_+$ ( $\sum_{i \in \mathcal{S}_+^{(M)}} \pi_i = 1$). This defines a proper probability distribution $\sum_{y \in \mathcal{S}^N} P_\theta(Y = y) = 1$ over all $S^N$ tip configurations (including extinct tips $Y_k = \varnothing$). Because $C(\varnothing \to \varnothing, \varnothing) = 1$ implies zero probability of any extant non-empty descendant arising from an $\varnothing$ ancestor, when evaluated with matched branch segmentation and uniform root weights over the $S_0 = |\mathcal{S}_+^{(M)}|$ allowed non-empty root states ($\pi_i = 1/S_0$, where $S_0 = S - 1$ on unconstrained single-layer trees), its log-likelihood on extant non-empty tips differs from unit-weight legacy pruning by the constant $-\ln S_0$:

   $$\ln L_{\text{unconditional\_complete}}(\theta) = \ln L_{\text{legacy, matched}}(\theta) - \ln S_0,$$

   yielding identical MLEs under matched segmentation while providing an exact complete-space forward simulator and probability-sum verification baseline (default legacy stratified runs use upstream decimal rounding on near-ultrametric branches, which can introduce minor parameter-dependent offsets relative to telescoping segmentation).

3. `tip_survival` : Retains rate-$e$ transitions from single-area states into $\varnothing$ and conditions the data probability on every sampled extant tip being non-empty ($Y_k \in \mathcal{S}_+$ for all $k = 1, \ldots, N$). Computing a second pruning pass $\tilde{\mathbf{L}}_{\text{root}}$ initialized with all-ones over allowed present-day non-empty tip states ($\tilde{L}_{k,i} = \mathbb{1}[i \in \mathcal{S}_+^{(1)}]$) yields the exact joint survival probability $P_{\text{surv}}(\theta) = \sum_{i \in \mathcal{S}_+^{(M)}} \pi_i^{(M)} \tilde{L}_{\text{root},i}$, and the conditioned log-likelihood:

   $$\ln L_{\text{tip\_survival}}(\theta) = \ln\left(\sum_{i \in \mathcal{S}_+^{(M)}} \pi_i^{(M)} L_{\text{root},i}\right) - \ln P_{\text{surv}}(\theta).$$

   Because observed non-empty tip data $Y$ already entail survival, for any latent range history $H$ at fixed parameters $\theta$, $P(H \mid Y, \text{surv}, T, \theta) = P(H \mid Y, T, \theta)$; tip-survival conditioning alters ancestral reconstructions and stochastic maps by shifting fitted parameter estimates $\widehat{\theta}$ and relative model weights.

**Extension to Multi-Stratum Time Slicing and Monotone Island Emergence:** Suppose geological time boundaries $0 = \tau_0 < \tau_1 < \ldots < \tau_M$ partition the phylogeny into strata $m = 1, \ldots, M$ from present to past, with allowed area sets $\mathcal{A}_1, \ldots, \mathcal{A}_M$ and induced allowed non-empty state sets $\mathcal{S}_+^{(m)} = \{s \in \mathcal{S}_+ : s \subseteq \mathcal{A}_m\}$. When islands or areas emerge chronologically from oldest to youngest ( $\mathcal{A}_M \subseteq \mathcal{A}_{M-1} \subseteq \ldots \subseteq \mathcal{A}_1$), every state $s \in \mathcal{S}_+^{(m)}$ active in an older stratum $m$ remains valid upon crossing boundary $\tau_{m-1}$ into younger stratum $m-1$ ($\mathcal{S}_+^{(m)} \subseteq \mathcal{S}_+^{(m-1)}$). Consequently: - Forward transitions across stratum boundaries preserve total probability mass without ad hoc projection or renormalization; - The root prior $\pi^{(M)}$ is normalized strictly over the non-empty states $\mathcal{S}_+^{(M)}$ permitted in the root's geological stratum; - Near-ultrametric trees (e.g., Hawaii *Psychotria*, tip-height spread $9.79 \times 10^{-4}$ Ma) are segmented via exact double-precision telescoping sub-branch lengths ( $\sum_k \Delta t_{e,k} = t_e$ to $10^{-16}$ precision), preventing artificial zero-length fossil-hook misclassification; - Conversely, non-monotone masks in which a previously allowed area disappears in a younger stratum ($\mathcal{A}_m \nsubseteq \mathcal{A}_{m-1}$) zero out probability mass across stratum boundaries without defining a physical extinction vs. range-contraction transition; `RustBEARS` explicitly rejects such masks under `tip_survival` / `unconditional_complete` while preserving them under `legacy_unconditioned` .

---

#### S1.3 Parameter Bounds, Zero-Boundary Re-Anchoring, and SHA-256 Checkpointing

- **Optimization Box Bounds**: Upstream compatibility templates retain historical positive floors ($d, e \in [10^{-12}, 5]$, cladogenetic jump relative weight $j \in [10^{-5}, 3 - 10^{-5}]$ for `DEC+J` , $[10^{-5}, 2 - 10^{-5}]$ for `DIVALIKE+J` , and $[10^{-5}, 1 - 10^{-5}]$ for `BAYAREALIKE+J` ), whereas explicit zero lower bounds ( `min = 0.0` ) are supported directly via `with_parameters(...)` and enforced automatically during nested `bootstrap_lrt` and zero-anchored profiling.
- **Exact Zero Probing & Re-Anchoring in `profile_ci`** : When historical templates supply a small numerical floor (e.g., $j_{\min} = 10^{-5}$), `profile_ci(..., eval_exact_zero = True, multi_basin_guard = True)` evaluates the restricted likelihood at exact physical `0.0` ; if $\ln L(0) \geq \ln L(\widehat{\theta})$, the reference optimum is re-anchored and `status_lower` is labeled `at_physical_zero` (distinguished from `infeasible_zero_approached` when $d = 0$ makes multi-area tips structurally impossible).

- **Tamper-Evident Checkpointing**: `bootstrap_lrt` and `fit_over_trees` serialize per-replicate JSONL/RDS checkpoints header-stamped with a canonical SHA-256 digest covering tree Newick topology/lengths, tip geography, stratum matrices, parameter bounds, conditioning convention, precision mode, seed, and native library version, rejecting mismatched resume attempts.

#### S2. Comprehensive Numerical Equivalence and Fair Runtime Benchmarks

##### Supplementary Table S2A. Archival Compatibility Across 77 Published Studies

| Verification Dimension | Cohort Size | Comparison Mode | Median Discrepancy | 95th Percentile | Maximum Discrepancy | Pass / Concordance Rate (Criterion) |
|---|---|---|---|---|---|---|
| Fixed-parameter $\lvert\Delta \ln L\rvert$ vs Original `BioGeoBEARS` (v1.1.4) | 1,339 configs (693 unstratified baseline + 646 stratified/modifier configs; 75 valid screens up to 638 states + 2 initial-screen timeouts recovered up to 848 states) | `LegacyF32` | $9.95 \times 10^{-14}$ | $1.59 \times 10^{-12}$ | $8.00 \times 10^{-11}$ | **100% (1,339 / 1,339)** ( $\lvert\Delta \ln L\rvert \le 10^{-9}$) |
| Root state marginal probability $\lvert\Delta P\rvert$ vs Original | 1,339 configs | `LegacyF32` | $3.33 \times 10^{-16}$ | $5.29 \times 10^{-16}$ | $4.11 \times 10^{-15}$ | **100% (1,339 / 1,339)** ( $\lvert\Delta P\rvert \le 10^{-12}$) |
| Internal node ancestral probability $\lvert\Delta P\rvert$ vs Original | 1,339 configs | `LegacyF32` | $6.66 \times 10^{-16}$ | $4.55 \times 10^{-15}$ | $1.50 \times 10^{-14}$ | **100% (1,339 / 1,339)** ( $\lvert\Delta P\rvert \le 10^{-12}$) |
| R vs Python dual-frontend relative $\lvert\Delta \ln L\rvert / \lvert\ln L\rvert$ | 77 studies | `LegacyF32` (matched precision) | $2.13 \times 10^{-16}$ | $9.86 \times 10^{-16}$ | $2.56 \times 10^{-15}$ | **100% (77 / 77)** (rel. $\lvert\Delta \ln L\rvert \le 10^{-12}$) |
| `StrictF64` vs `LegacyF32` fixed-parameter $\lvert\Delta \ln L\rvert$ | 1,339 configs | Cross-precision | $1.41 \times 10^{-6}$ | $2.82 \times 10^{-5}$ | $3.62 \times 10^{-4}$ | `StrictF64` vs `LegacyF32` root argmax **99.86% (692 / 693)** (vs Original R **98.70% [684 / 693]**) |
| Independent cold-start MLE non-inferiority vs Original | 693 baseline fits | `LegacyF32` (matched precision) | $+7.63 \times 10^{-7}$ | $+5.27 \times 10^{-2}$ | +17.43 lnL gain | **100% non-inferior (693 / 693)** ($\Delta \ln L \ge -10^{-3}$; worst $-8.07 \times 10^{-5}$) |
| Paired whole-tree Uniformization vs Original BSM event TVD | 30 studies × 200 maps | Cross-engine | 0.0019 (TVD) | 0.0071 (TVD) | 0.1034 (TVD; route $r = 0.9973$ mean, 0.9995 median) | 12 / 12 exact match in `legacy_r_bridge` |

#### Supplementary Table S2B. Equal-Core (1 vs 1), Equal-Resource (8 vs 8), and Best-Baseline Runtime Benchmarks on Idle AMD EPYC 7763

*(All timings are medians across repeated runs on an idle host [* `loadavg ≈ 0.3–3.1` *]. For Original* `BioGeoBEARS` *8-core BSM, 200 maps were parallelized across 8* `parallel::mclapply` *worker processes [8 × 25 maps]; for Original 8-core MLE,* `num_cores_to_use = 8` *was passed to* `bears_optim_run`*, where upstream SOCK cluster serialization overhead degrades wall-clock speed relative to 1 core on 64–128 state spaces and all 4-stratum runs, while slightly reducing unstratified 256-state runtime from 115.44 s to 98.97 s. Both 8-vs-Best [8-core* `RustBEARS` *vs* `min(1-core, 8-core)` *Original* `BioGeoBEARS`*, plotted in Figure 1C] and strict 8-vs-8 speedups are reported below.)*

| Task & Dataset Configuration | Original R (1 Core) | Original R (8 Cores) | `RustBEARS` R (1 Thread) | `RustBEARS` R (8 Threads) | `RustBEARS` Python (1 Thread) | `RustBEARS` Python (8 Threads) | 1-vs-1 Speedup (R / Py) | 8-vs-Best Orig. (R / Py) | 8-vs-8 Speedup (R / Py) |
|---|---|---|---|---|---|---|---|---|---|
| **BSM (200 maps)**: *Psychotria* single-layer (16 states) | 12.59 s | 1.18 s | 1.29 s* | 1.21 s* | 0.014 s | 0.013 s | **9.8× / 907×** | **1.0×* / 91.2×** | **1.0×* / 91.2×** |
| **BSM (200 maps)**: *Psychotria* M3 4-stratum (16 states) | 101.28 s | 5.55 s† | 1.65 s* | 1.72 s* | 0.072 s | 0.026 s | **61.2× / 1,405×** | **3.2×* / 212.7×** | **3.2×* / 212.7×** |
| **BSM (200 maps)**: Synthetic 64 states (single-layer) | 35.18 s | 3.06 s | 2.75 s* | 2.32 s* | 0.514 s | 0.084 s | **12.8× / 68.4×** | **1.3×* / 36.6×** | **1.3×* / 36.6×** |
| **BSM (200 maps)**: Synthetic 128 states (single-layer) | 47.98 s | 5.12 s | 3.65 s* | 2.53 s* | 1.267 s | 0.197 s | **13.1× / 37.9×** | **2.0×* / 25.9×** | **2.0×* / 25.9×** |
| **MLE (** `DEC+J` **)**: 64 states (single-layer, 20 tips) | 7.87 s | 108.60 s | 2.79 s | 1.92 s | 1.12 s | 0.97 s | **2.8× / 7.0×** | **4.1× / 8.1×** | **56.6× / 111.7×** |
| **MLE (** `DEC+J` **)**: 128 states (single-layer, 20 tips) | 29.59 s | 78.39 s | 6.02 s | 2.75 s | 2.51 s | 1.93 s | **4.9× / 11.8×** | **10.8× / 15.4×** | **28.5× / 40.7×** |
| **MLE (** `DEC+J` **)**: 256 states (single-layer, 20 tips) | 115.44 s | 98.97 s | 8.63 s | 4.16 s | 12.76 s | 9.09 s | **13.4× / 9.0×** | **23.8× / 10.9×** | **23.8× / 10.9×** |
| **MLE (** `DEC+J` **)**: 64 states (4 strata, 20 tips) | 33.88 s | 272.57 s | 4.05 s | 2.91 s | 2.73 s | 1.33 s | **8.4× / 12.4×** | **11.7× / 25.5×** | **93.8× / 205.5×** |
| **MLE (** `DEC+J` **)**: 128 states (4 strata, 20 tips) | 73.51 s | 162.96 s | 7.75 s | 4.23 s | 6.27 s | 3.58 s | **9.5× / 11.7×** | **17.4× / 20.5×** | **38.6× / 45.5×** |

| Task & Dataset Configuration | Original R (1 Core) | Original R (8 Cores) | RustBEARS R (1 Thread) | RustBEARS R (8 Threads) | RustBEARS Python (1 Thread) | RustBEARS Python (8 Threads) | 1-vs-1 Speedup (R / Py) | 8-vs-Best Orig. (R / Py) | 8-vs-8 Speedup (R / Py) |
|---|---|---|---|---|---|---|---|---|---|
| **MLE ( DEC+J ):** 256 states (4 strata, 20 tips) | 187.83 s | 261.52 s | 15.57 s | 6.83 s | 10.85 s | 5.21 s | **12.1× / 17.3×** | **27.5× / 36.1×** | **38.3× / 50.2×** |

**Note on R `runBSM` vs Python `run_bsm`: Both interfaces return complete per-event cladogenetic/anagenetic histories and route summary tables; a single top-level R `runBSM` call additionally converts upstream S4/list structures into legacy R `RES_clado_events_tables` and `RES_ana_events_tables` lists (~1.1 s fixed R-side formatting cost), which dominates small 200-map batches but amortizes on 1,000–5,000 maps (e.g., 5,000 stratified maps in ~2.2 s). †In Original `BioGeoBEARS` M3 stratified 8-core `mclapply`, occasional worker chunks encounter upstream `events_txt_list_into_events_table` decimal sub-edge matching errors on near-ultrametric tips; `RustBEARS` completes 100% of maps without error.*

---

#### Supplementary Table S2C. Full Six-Stage Uncertainty Workflow Wall-Clock Time and Peak RSS Across State-Space Sizes

| State Space (Regions) | model_average (6 models, 8T) | profile_ci ($\chi^2_1$, 8T) | profile_ci (plug-in boot., 8T) | bootstrap_lrt ( $B = 199$, 8T) | run_bsm (5,000 maps, 8T) | fit_over_trees (20 trees, 8T) | Process Peak RSS (8T) | Scheduling & Single-Thread Notes |
|---|---|---|---|---|---|---|---|---|
| **16 states** ($r = 4$) | 2.07 s | 14.04 s | 170.13 s | 186.90 s | 1.47 s | 39.02 s | **161–232 MiB** | Exclusive isolated host; all stages complete at 1T ( `4.7–925 s` ) |
| **64 states** ($r = 6$) | 8.55 s | 67.27 s | 738.67 s | 605.73 s | 4.75 s | 125.25 s | **162–232 MiB** | Exclusive isolated host; all stages complete at 1T ( `28.9–5,440 s` ) |
| **256 states** ($r = 8$) | 35.12 s* | 482.59 s* | 4,838.86 s* | 3,938.94 s* | 30.10 s* | 741.22 s* | **168–265 MiB** | *Batch-scheduled wall times; at 1T, `profile_plugin` & `bootstrap_lrt` hit 14,400 s (4 h) cap |

*(Note: When retaining 10,000 full BSM event-history tables simultaneously in memory during the 200-tree Trimezieae multi-tree run, Python process Peak RSS measured 1,703 MiB owing to retained event arrays rather than kernel workspace.)*

---

#### S3. Statistical Calibration and Archival Tip-Survival Reassessment Details

##### Supplementary Table S3A. Null `+J` ($j = 0$) Extinction Recovery and Spurious `+J` Selection Across Fixed-Tree ($3 \times 200$) and Independent Birth–Death Pruned-Tree ($3 \times 200$) Simulations

*(Note: In the birth–death pruned-tree simulations, phylogenies and ranges were simulated forward in continuous time via an independent Gillespie birth–death + standard DEC range-evolution generator [expansion rate $|A| \cdot d$ per unoccupied destination area, stopping uniformly in the first sojourn at $N = 30$ extant lineages with $\geq 2$ represented areas; Hartmann et al. 2010] where losing the final occupied area causes lineage extinction and extinct side lineages are pruned from the reconstructed tree [$r = 4$ regions, 16 states]. Nested `DEC+J` fits include null-optimum warm starts [0 negative D values beyond numerical tolerance, minimum raw $D = -8.5 \times 10^{-14}$]. Both median and mean/RMSE of $\widehat{e}$ are reported because single-tree $\widehat{e}$ distributions exhibit zero-boundary point mass plus right skewness: in the $e = 0.15$ pruned-tree regime, conditioning lowers median absolute error from $0.150$ to $0.111$ [closer to truth on 125/200 trees, farther on 59/200, tied on 16/200], whereas unconditioned collapse to $\widehat{e} = 0$ produces an artificial RMSE equal to $e_{\text{true}}$.)*

| Simulation Regime ( $j_{\text{true}} = 0$) | Method / Convention | $\widehat{e}$ Median [IQR] | $\widehat{e}$ Mean (RMSE) | $\widehat{e} \approx 0$ Frac. | $\widehat{e} = 5.0$ Cens. | Exact $D = 0$ Frac. | AICc `+J` FP Rate | $\bar{\chi}^2_{0.5}$ FP Rate | $\chi^2_1$ FP Rate |
|---|---|---|---|---|---|---|---|---|---|
| **Fixed tree**: $d = 0.015, e = 0.02$ | `legacy_unconditioned` | 0.000 [0.000, 0.000] | 0.000 (0.020) | 100.0% | 0.0% | 64.5% | 5.5% | 5.0% | 3.0% |
| | `DEC*` (no 1-area ext.) | 0.005 [0.000, 0.144] | 0.200 (0.665) | 46.0% | 1.0% | 76.5% | 1.5% | 1.5% | 0.0% |
| | **`tip_survival`** | **0.026** [0.000, 0.154] | 0.272 (0.945) | 42.5% | 2.5% | 77.5% | **0.5%** | **0.5%** | **0.0%** |
| **Fixed tree**: $d = 0.015, e = 0.08$ | `legacy_unconditioned` | 0.000 [0.000, 0.000] | 0.000 (0.080) | 100.0% | 0.0% | 44.0% | **18.0%** | **16.5%** | 8.5% |
| | `DEC*` (no 1-area ext.) | 0.062 [0.000, 0.236] | 0.275 (0.670) | 34.0% | 1.0% | 63.0% | 4.5% | 4.0% | 1.0% |
| | **`tip_survival`** | **0.093** [0.000, 0.321] | 0.359 (0.917) | 36.0% | 2.0% | 64.5% | **1.5%** | **1.0%** | **0.5%** |
| **Fixed tree**: $d = 0.015, e = 0.15$ | `legacy_unconditioned` | 0.000 [0.000, 0.000] | 0.000 (0.150) | 100.0% | 0.0% | 35.0% | **22.0%** | **21.0%** | 14.0% |
| | `DEC*` (no 1-area ext.) | 0.120 [0.004, 0.375] | 0.317 (0.604) | 23.0% | 0.5% | 60.5% | 2.5% | 2.0% | 1.5% |
| | **`tip_survival`** | **0.145** [0.008, 0.433] | 0.377 (0.766) | 23.0% | 1.0% | 65.5% | **1.5%** | **1.5%** | **1.0%** |
| **BD Pruned**: $\lambda = 0.30, d = 0.03, e = 0.08$ | `legacy_unconditioned` | 0.000 [0.000, 0.000] | 0.001 (0.079) | 90.0% | 0.0% | 39.0% | **21.0%** | **19.5%** | 13.5% |
| | `DEC*` (no 1-area ext.) | 0.076 [0.008, 0.195] | 0.205 (0.596) | 22.5% | 1.0% | 64.0% | 4.5% | 4.5% | 2.0% |

| Simulation Regime ( $j_{\text{true}} = 0$) | Method / Convention | $e$ Median [IQR] | $e$ Mean (RMSE) | $e \approx 0$ Frac. | $e = 5.0$ Cens. | Exact $D = 0$ Frac. | AICc +J FP Rate | $\chi^2_{0.5}$ FP Rate | $\chi^2_1$ FP Rate |
|---|---|---|---|---|---|---|---|---|---|
| | **`tip_survival`** | **0.079** [0.011, 0.224] | 0.203 (0.575) | 22.5% | 1.0% | 66.0% | **4.5%** | **4.0%** | **1.5%** |
| **BD Pruned**: $\lambda = 0.35, d = 0.04, e = 0.15$ | `legacy_unconditioned` | 0.000 [0.000, 0.000] | 0.001 (0.149) | 84.5% | 0.0% | 25.5% | **33.0%** | **31.5%** | **22.0%** |
| | `DEC*` (no 1-area ext.) | 0.175 [0.070, 0.343] | 0.268 (0.423) | 8.0% | 0.0% | 59.0% | 4.0% | 3.0% | 1.0% |
| | **`tip_survival`** | **0.183** [0.094, 0.365] | 0.266 (0.323) | 8.0% | 0.0% | 62.5% | **4.0%** | **2.5%** | **1.0%** |
| **`DEC*`-Truth Control**: $\lambda = 0.30, d = 0.03, e = 0.10$ | `legacy_unconditioned` | 0.000 [0.000, 0.000] | 0.001 (0.099) | 92.0% | 0.0% | 37.5% | **20.5%** | **19.5%** | 13.0% |
| | `DEC*` (true model) | 0.079 [0.000, 0.225] | 0.207 (0.478) | 28.5% | 0.5% | 66.0% | 5.5% | 5.0% | 1.0% |
| | **`tip_survival`** | **0.101** [0.000, 0.212] | 0.196 (0.438) | 31.0% | 0.5% | 66.0% | **5.0%** | **4.5%** | **1.0%** |

---

#### Supplementary Table S3B. Summary of Single-Layer and Multi-Stratum Uncertainty Module Calibrations

| Validation Module & Design | Replicates | Key Calibration Result | Boundary / Censoring Behavior |
|---|---|---|---|
| **Profile Likelihood Intervals** (40 tips, 16 states, 4 scenarios: interior, $e = 0$, $j = 0$, and $e = j = 0$) | 800 datasets | Asymptotic $\chi^2_1$ `profile_ci` 100% available: interior $d, j$ coverage **94.0%–98.5%**, interior/boundary $e$ coverage **99.5%–100.0%**; boundary $j = 0$ coverage **99.5%–100.0%** | Central-difference Hessian precheck fails in **54%–91%** of fits at box boundaries (57%–91% across the scenarios plotted in Fig. 2B); $e$ upper bound right-censored at 5.0 in **18.5%–83.0%** of 40-tip datasets |
| **Four-Stratum Chronological Island Emergence** (30 tips, 4 islands `A -> AB -> ABC -> ABCD`, $d = 0.08, e = 0.06, j = 0$) | 100 datasets ($B = 99$) | Median $\widehat{d} = 0.082, \widehat{e} = 0.028$; stratified `profile_ci` coverage: $d = 95.0\%$ ( `[88.7%, 98.4%]` ), $e = 99.0\%$ ( `[94.6%, 100.0%]` ); stratified `bootstrap_lrt` Type-I error **3.0%** ( `[0.6%, 8.5%]` ) | **0.0%** upper-box censoring on both $d$ and $e$; asymptotic $\chi^2_1$ / $\bar{\chi}^2_{0.5}$ conservative ( `0.0%` ) |

| Validation Module & Design | Replicates | Key Calibration Result | Boundary / Censoring Behavior |
|---|---|---|---|
| **Parametric Bootstrap LRT** (30 tips, 16 states, null $j = 0, e = 0.02$ vs weak alternative $j = 0.03, e = 0.02, B = 99$) | 400 datasets | Under `tip_survival`: null Type-I error **4.0%** ( `[1.7%, 7.7%]` ) vs $\chi^2_1$ `0.0%` / $\bar{\chi}^2_{0.5}$ `1.5%`; weak $j = 0.03$ power **11.0%** ( `[7.0%, 16.2%]` ) vs $\chi^2_1$ `1.5%` | Scoring `legacy_unconditioned` test statistics while overriding bootstrap generation with `tip_survival` gives 5.0% null rejection at $e = 0.02$ and 69.0% ( `138/200` ) rejection at $j = 0.03$, $e = 0.02$; however, because unconditioned null fits collapse $\widehat{e}_{\text{null}} \to 0$ under moderate-to-high extinction, self-consistent `tip_survival` fitting and bootstrap generation is recommended for inference |
| **Akaike Multimodel Averaging** (30 tips, 16 states, comparable-weight candidate regimes) | 600 datasets | Mean ancestral node Brier reduction vs single best model: **`-0.048`** ( `[-0.056, -0.041]` ) and **`-0.022`** ( `[-0.027, -0.017]` ) | Eliminates $+\infty$ log-loss ($-\infty$ log-score) incurred by single-model selection in up to **21.0%** ( `63/300` ) of datasets |
| **Stochastic Mapping MCSE & Tree-Chain ESS Gate** ( `fixed_sample`, `independent`, and `mcmc_chain` at $\rho = 0.8$) | 3,400 runs | Fixed-sample coverage **92.0%–95.8%**; independent $K = 10$ ($t_9$) **95.2%**; long MCMC ( $K = 800$, median $\widehat{\text{ESS}} = 107$) passes gate in **99.0%–99.2%** with **93.5%–93.7%** conditional coverage | Short MCMC ($K = 60$, true ESS `8.4`) blocked in **99.8% ( `499/500` )** of runs (unguarded normal intervals yield **79.6%–79.8%** coverage) |

*(Dataset total across single-tree and multi-model simulation experiments:* $600$ [fixed-tree null] $+ 400$ [birth–death pruned] $+ 200$ [DEC* control] $+ 800$ [profile CI] $+ 100$ [four-stratum] $+ 400$ [bootstrap LRT] $+ 600$ [model averaging] $= 3,100$ *simulated datasets.)*

---

#### Supplementary Table S3C. Empirical Reassessment of Tip-Survival Conditioning Across the 77-Study Archival Corpus

| Corpus Stratum / Metric | Count / Value | Biological & Statistical Interpretation |
|---|---|---|
| Total published studies in archival benchmark | 77 studies | 75 valid baseline tracks (2 upstream timeout-skipped baselines: `BSM129`, `BSM143` ) |
| Explicitly rejected outside `tip_survival` contract | 15 studies | 10 contain non-contemporaneous fossil tips ( $\geq 1$ Ma); 4 extant trees exhibit non-ultrametric tip offsets ($0.004$–$0.401$ Ma); 1 triggers numerical survival underflow ( `BSM161` ) |
| Skipped above benchmark state cap ( $> 512$ states) | 2 studies | High-area datasets ( `BSM018` : 638 states; `BSM043` : 562 states) retained in `legacy_unconditioned` |
| **Evaluated extant archival entries under `tip_survival` ( `DEC & DEC+J` )** | **58 entries** | Standardized baseline comparison with null-optimum warm starts ( `BSM050`, `BSM127` verified non-negative $D$) |
| `legacy_unconditioned` `DEC` near-zero extinction ($\widehat{e} \leq 10^{-4}$) | **36 / 58 (62.1%)** | Median $\widehat{e} = 0.000$ across archival entries |

| Corpus Stratum / Metric | Count / Value | Biological & Statistical Interpretation |
|---|---|---|
| `tip_survival` `DEC` finite interior point estimates ($0 < \widehat{e} < 5.0$) | **51 / 58 (87.9%)** | **0 / 58 (0.0%)** near zero (minimum $\widehat{e} = 0.0028$); median interior $\widehat{e} = 0.110$ (overall median `0.186`); an interior MLE does not guarantee a finite 95% profile upper bound (`BSM169` [*Osmia*, 32 tips, $\widehat{e} = 0.95$] yields closed CI `[0.23, 4.03]`, whereas small/high-$\widehat{e}$ `BSM039` [6 tips] and `BSM058` [$\widehat{e} = 4.93$] remain right-censored at 5.0) |
| `tip_survival` `DEC` box-upper-bound censoring ($\widehat{e} = 5.0$) | **7 / 58 (12.1%)** | Accompanied by extreme survival log-probabilities ($\ln P_{surv} = -325.5$ to $-6,419.3$) and flat upper-tail likelihood ridges; all **4 / 4 (100%)** archival entries with 100% single-area-coded tips hit 5.0 (censored median **100.0%**, range **70.0%–100.0%**, vs **83.3%** in interior entries; `DEC+J` hits 5.0 in **3 / 58 [5.2%]**) |
| Archival entries where AICc prefers `DEC+J` over `DEC` (all 58 clades) | **44 / 58 (75.9%)** → **21 / 58 (36.2%)** | 23 entries shift from `DEC+J` to `DEC` and 0 reverse (17 of 21 remaining `DEC+J` preferences have $\Delta\text{AICc} \geq 2$); median LRT $D$ drops from **18.70** to **0.64** |
| Sensitivity check: AICc `DEC+J` preference excluding 7 ceiling-censored clades | **37 / 51 (72.5%)** → **20 / 51 (39.2%)** | Confirms that reduced `DEC+J` preference is not an artifact of ceiling-censored `DEC` fits |

*(Ceiling sensitivity note on stratified Hawaiian Psychotria [5,000 six-model-averaged maps per ceiling]: varying* $e_{max} \in \{1, 5, 20\}$ *preserves progressive older-to-younger colonization as the dominant flux [**88.3%, 81.3%, and 78.4%** of all transitions, with* K→O *consistently ranked first at 4.06, 5.99, and 7.08], whereas secondary route magnitudes and ordering between* O→M *[2.55, 2.93, 3.34] and* K→M *[1.26, 2.87, 3.63] shift with the extinction ceiling.)*

---

#### S4. External Bayesian Phylogenetics (`RevBayes`) Posterior Tree Propagation Study

To evaluate `fit_over_trees` on trees inferred externally from molecular alignments rather than fixed-topology simulations, we simulated 10 datasets (50 extant tips, 2,000 bp JC69 alignments under a coalescent tree prior + discrete biogeography; all 10 passed MCMC convergence diagnostics and 9 completed primary biogeographic scoring under default branch-length thresholds) and inferred posterior tree samples in `RevBayes` v1.4.2 (two independent MCMC chains per dataset, 200 thinned posterior trees per dataset) compared against single Maximum Clade Credibility (MCC) tree inference:

1. **Clade-Conditional Accuracy Across Support Strata**: Across all 399 clades shared between the MCC tree and the posterior tree sample (9 scored datasets), mean ancestral Brier score was nearly identical between multi-tree propagation (0.3106) and single-MCC-tree inference (0.3124; paired difference multi-tree minus MCC $= -0.0018$, dataset-level 95% bootstrap CI $[-0.0160, +0.0112]$, deterministic 10-dataset missing-data bounds $[-0.2016, +0.1984]$). Stratified by posterior clade support $P_{clade}$, paired multi-tree minus MCC Brier differences were $-0.0031$ ($[-0.0164, +0.0088]$ across 370 high-support clades with $P_{clade} > 0.95$; $-0.0020$ across the 337 universally supported clades with $P_{clade} = 1.0$), $+0.0243$ ($[+0.0008, +0.0498]$ across 13 moderate-support clades with $0.70 \leq P_{clade} \leq 0.95$), and $+0.0134$ ($[-0.0061, +0.0458]$ across 16 low-support clades with $P_{clade} < 0.70$, where MCC had lower Brier error on 6 of 8 datasets). Thus, conditioning on weakly supported alternative trees does not systematically lower raw node-wise Brier error relative to a single summary tree; rather, the inferential role of `fit_over_trees` is to prevent weakly supported clades ($P_{clade} < 1$) from being reported as topologically certain, separating topological monophyly support ($P_{clade}$) from clade-conditional ancestral range probabilities and reporting deterministic bounds $[P_{min}, P_{max}]$ across any missing tree fits.

2. **Coalescent Near-Zero Branch Guard**: In the remaining replicate (`dataset_008`), rapid speciation produced an internal branch of length $5.34 \times 10^{-7}$, triggering initial rejection under the default `min_branchlength = 1e−6` threshold; lowering `min_branchlength = 1e−9` completed all 200/200 posterior trees cleanly, motivating an explicit check on minimum positive branch lengths when importing coalescent posterior tree sets.
